# Ataxia-Telangiectasia Mutated Inhibition Enhances Adeno-Associated Virus-Mediated Knock-in Efficiency in Pig Zygotes

**DOI:** 10.64898/2025.12.20.695642

**Authors:** Hiromasa Hara, Mitsuhiro Noguchi, Fuminori Tanihara, Makoto Inoue, Yutaka Hanazono, Arata Honda

**Author notes:** H.H. and A.H. served as co-corresponding authors. Arata Honda, PhD; **Email:** Hiromasa Hara, PhD; **Email:**. H.H and M.N contributed equally to this work.

## Abstract

A persistent global shortage of transplantable organs has renewed interest in pigs as donors for xenotransplantation and hosts for interspecies organogenesis considering their physiology closely resembles that of humans. Such applications require pigs harboring genetic modifications, including knockouts, human transgenes, and conditional alleles. Although such complex allele combinations can be assembled by sequential editing of fetal fibroblasts, followed by somatic cell nuclear transfer (SCNT), SCNT is often associated with developmental abnormalities. Direct genome editing of pig zygotes offers an alternative route; however, achieving long-fragment knock-in, particularly at multiple loci, remains challenging. Here, adeno-associated virus (AAV) donor vectors for long-fragment knock-in in pig zygotes were evaluated, and tested whether transient inhibition of ataxia-telangiectasia mutated (ATM), which becomes overactivated during AAV transduction, could enhance knock-in efficiency. AAV6 donors supported efficient ACTB–mEGFP knock-in (33%), whereas short-term treatment with the ATM inhibitor AZ32 increased the efficiency to 66%. In addition, AZ32 enhanced simultaneous double knock-in at *ACTB* and *UBB*, generating a higher proportion of double-positive blastocysts (52% vs. 19% in controls), without affecting blastocyst formation rates or total cell numbers. Embryo transfer yielded ACTB–mEGFP and UBB–mScarlet-I3 double knock-in fetuses (33%), including an F0 fetus composed almost entirely of double knock-in cells. These results demonstrate that combining AAV donors with transient ATM inhibition is an efficient strategy for long-fragment knock-in at multiple loci in pigs.

## Introduction

Pigs are considered one of the most promising donor species for clinical xenotransplantation because their physiology resembles that of humans (1, 2). Recent first-in-human studies using genetically engineered donor pigs for heart, kidney, and liver transplantation have highlighted the rapid progress in the field and the need for donor animals that carry complex combinations of genetic modifications to regulate immunological, coagulation, and growth pathways (3–6). In addition, pigs are increasingly being positioned as hosts for interspecies organogenesis in which genetically defined organ deficiencies in the host are complemented by human pluripotent stem cells to produce human-compatible organs or tissues (7–11). The emerging applications require pigs that harbor several defined genetic modifications, including gene knockouts, human transgenes, and conditional alleles that allow temporal or lineage-restricted ablation of specific organs or cell populations (12, 13).

To date, all pigs carrying such complex allele combinations have been produced by sequentially editing fetal fibroblasts *in vitro,* followed by somatic cell nuclear transfer (SCNT). SCNT allows multiple targeted modifications to be accumulated in a single cell and has therefore become the dominant method for producing donor lines for xenotransplantation (14). In particular, a recent study that produced pigs carrying 69 genetic modifications (15) clearly demonstrated this advantage, highlighting the powerful capacity of SCNT for application in large-scale genome engineering. However, SCNT requires specialized technical skills and incomplete nuclear reprogramming can cause epigenetic abnormalities that impair survival during prenatal and postnatal development (16–19), thereby limiting the direct use of SCNT-derived pigs for research or clinical applications. Although such abnormalities can be overcome by passing the genome through the germline, such as by natural breeding (18, 20, 21), this approach requires the generation of both male and female SCNT-derived lines, making the process logistically demanding and adding considerable time and cost requirements.

Direct delivery of CRISPR/Cas9 into pig zygotes by microinjection or electroporation has emerged as a key alternative to SCNT and is widely used currently to generate gene knock-outs (22–25). This zygote-based approach has also been used to introduce point mutations or short sequences using single-stranded oligodeoxynucleotide donors (26–28). However, achieving efficient long-fragment knock-in, even at a single locus, remains technically challenging. The only report of long-fragment knock-in in pig zygotes, generated by microinjection of Cas9 together with a plasmid donor, targeted a locus at which gene disruption is lethal. Consequently, embryos lacking a correctly integrated donor allele would be eliminated via strong negative selection during development (29). In mice and rats, this limitation has been addressed by combining Cas9 ribonucleoprotein (RNP) with adeno-associated virus (AAV) donor vectors, an approach that has so far been established only in rodent zygotes (30–32). However, as AAV-based strategies have not yet achieved sufficient efficiency to support simultaneous long-fragment knock-in at multiple loci, rodent studies typically establish single-locus knock-in lines independently and rely on subsequent intercrossing to assemble multiple alleles (31, 33). In pigs, by contrast, AAV-mediated long-fragment knock-in in zygotes has not yet been established, and the rodent-style approach of generating multiple single-locus knock-in lines for subsequent intercrossing is impractical because of the substantial time, cost, and large-animal housing resources required.

In a recent work, the authors of the present study observed that linear DNA derived from AAV activates the ataxia-telangiectasia mutated (ATM) kinase in a dose–dependent manner, which is a central regulator of the cellular DNA damage response (Natsagdorj et al., *Commun Biol* in Revision). Excessive ATM activation induces apoptosis in cells transduced with high copies of AAV genomes, even though the cells are otherwise capable of supporting high knock-in efficiencies, and such activation also increases classical non-homologous end-joining (c-NHEJ), a repair pathway that competes directly with knock-in. Building on such observations, the authors of the present study further showed that short-term pharmacological inhibition of ATM increases AAV-mediated knock-in efficiencies markedly across multiple cell types, including mouse pluripotent stem cells, human cell lines, and mouse hematopoietic stem cells (Natsagdorj et al., *Commun Biol* in Revision) (34).

In the present study, AAV donor vectors in pig zygotes are tested for the first time to evaluate their capacity to support long-fragment knock-in. Furthermore, whether transient inhibition of ATM enhances AAV-mediated knock-in efficiency, including simultaneous multiple knock-ins, is examined.

## Results

### Establishment of a High Signal-to-Noise Knock-in Reporter System and Initial Screening of ATM Inhibitors Using Immortalized Pig Mesenchymal Stromal Cells

To enable the systematic evaluation of long-fragment knock-ins in pig cells, the authors first established an immortalized cell line derived from pigs with the same genetic background as that used for gamete collection (*SI Appendix*, Fig. S1). Primary adherent cells derived from pig bone marrow displayed limited proliferative capacity, showing progressively slower growth over early passages and eventually failing to continue to expand. To overcome the limitation, electroporation of a plasmid encoding the SV40 large T antigen, followed by puromycin selection, conferred stable long-term proliferation while maintaining stromal cell-like morphology (*SI Appendix*, Fig. S1A and B). Subsequently, whether the immortalized cells exhibited mesenchymal stromal cell (MSC)-like properties was assessed by analyzing their surface marker expression levels. Cells expressing high levels of CD29, CD44, CD105, and CD166 essentially exhibited no CD45 expression, and exhibited dim but reproducible CD73 expression (*SI Appendix*, Fig. S1C). The marker profile is consistent with previously described pig primary MSCs, which can undergo adipogenic, osteogenic, and chondrogenic differentiation (35). The results confirmed successful establishment of an immortalized pig MSC line with a genetic background matching that of the pigs used for gamete collection.

A knock-in reporter system for pigs was established using immortalized porcine MSCs (Fig. 1A). The system uses an AAV donor carrying an mEGFP cassette flanked by 1.5-kb homology arms designed to insert mEGFP in the frame at the C-terminus of the pig *ACTB* locus. To confirm that mEGFP expression accurately reflected targeted knock-in, three conditions in immortalized pig MSCs were tested, including Cas9 RNP alone, AAV donor alone, or both together (Fig. 1B). Flow cytometry showed no detectable mEGFP-positive cells under the RNP only condition, whereas AAV donor alone yielded a very minor mEGFP-positive fraction (0.01%). Notably, the population exhibited clearly elevated fluorescence intensity rather than diffuse background, which may reflect occurrence of a small number of knock-in events, even without Cas9-induced double-strand breaks (DSBs). In contrast, the RNP and AAV combination produced a clear mEGFP-positive population (12.22%), indicating that mEGFP expression depends predominantly on the presence of both genomic DSBs and the donor vector. The mEGFP-positive population exhibited fluorescence intensities that were approximately 300-fold higher than those observed in the negative fraction. PCR analysis of the sorted mEGFP-positive and mEGFP-negative fractions further demonstrated that knock-in-specific bands were detected only in the positive fraction, whereas the negative fraction lacked these junctions (Fig. 1C). Overall, the results demonstrate that the system can detect precise long-fragment knock-in events in pig cells with a very high signal-to-noise ratio.

**Figure 1.**
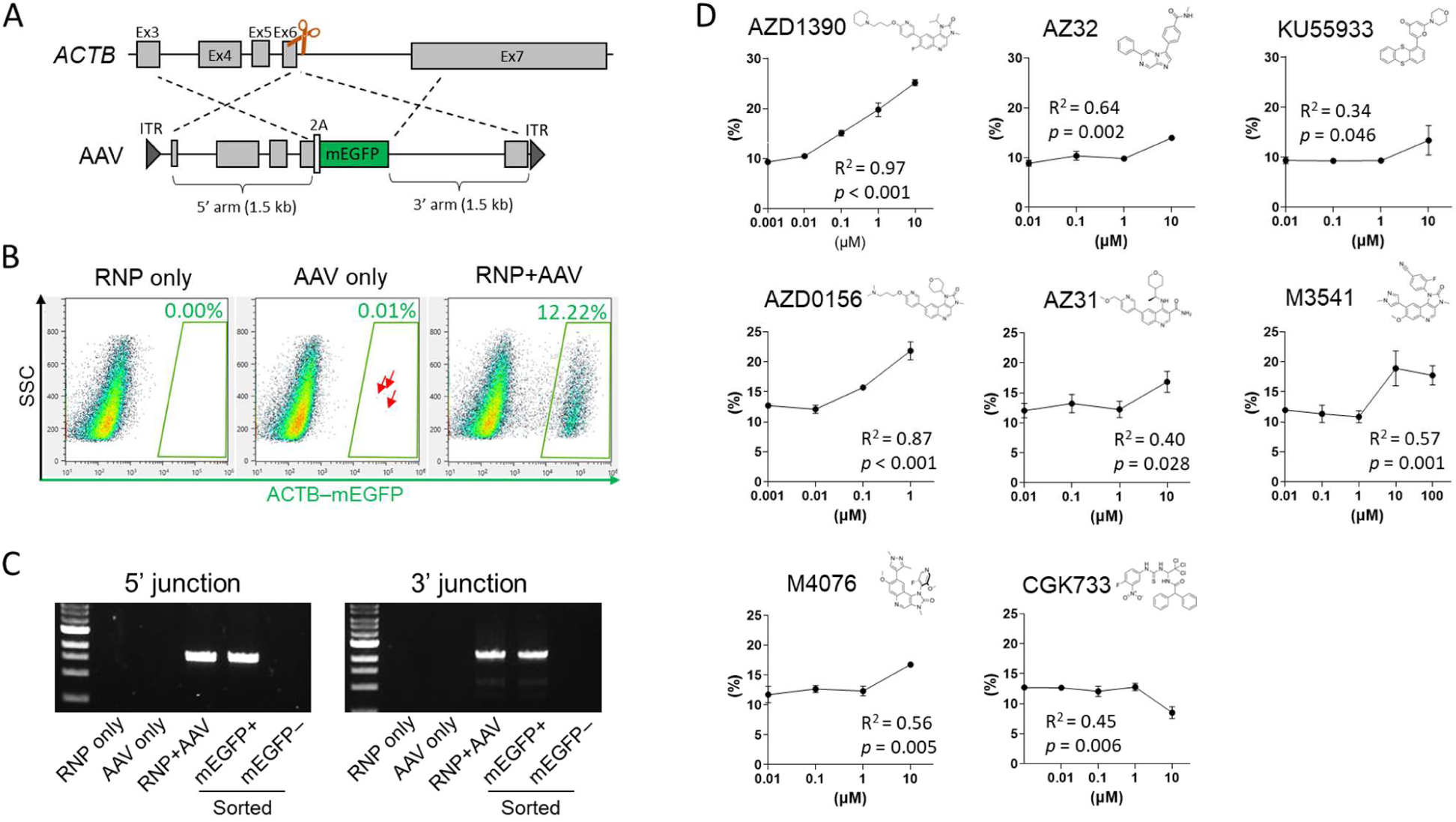
Establishment of a knock-in reporter system for pigs and screening of ataxia-telangiectasia mutated (ATM) inhibitors in immortalized pig mesenchymal stromal cells (MSCs). (*A*) Schematic of the ACTB–mEGFP knock-in strategy. An adeno-associated virus (AAV) donor carrying an mEGFP cassette linked via a 2A sequence and flanked by 1.5-kb homology arms was designed to insert mEGFP in frame at the C terminus of the pig ACTB. The orange scissors indicate the Cas9 cleavage site in exon 6. ITR, inverted terminal repeat. (*B*) Representative flow cytometry plots of immortalized pig MSCs 72 h after treatment with Cas9 RNP alone (RNP only), AAV donor alone (AAV only), or both together (RNP + AAV). The green gate denotes mEGFP-positive knock-in cells. Arrows indicate rare mEGFP-positive cells observed in the absence of Cas9 RNP. (*C*) Junction PCR analysis confirming correct 5’ and 3’ knock-in junctions using genomic DNA. (*D*) Dose–response curves for AAV-mediated knock-in in pig MSCs treated with the indicated ATM inhibitors. Regression lines represent log-linear fits of inhibitor concentration versus knock-in frequency, and statistical significance was evaluated by testing whether the regression slope differed from zero (n = 3 biological replicates).

Having established the reporter system, a panel of ATM inhibitors in the immortalized porcine MSCs was screened to assess their ability to promote AAV-mediated knock-in (Fig. 1D). Most compounds produced clear dose-dependent increases in the proportion of mEGFP-positive cells, with significant log-linear regression fits. In contrast, CGK733 failed to enhance knock-in and instead produced a modest yet significant decrease in knock-in efficiency at higher concentrations. Thus, CGK733 was not suitable for subsequent experiments in pig zygotes. The results indicate that the pharmacological inhibition of ATM represents a promising strategy for enhancing AAV-mediated knock-in in pig cells.

### Enhancement of Knock-in Efficiencies in Pig Zygotes by ATM Inhibitors

Building on the results obtained with immortalized pig MSCs, whether ATM inhibitors also enhance knock-in efficiency in pig zygotes was examined. Before performing inhibitor screening, the electroporation of Cas9 RNP into one-cell zygotes efficiently induced DSBs at the *ACTB* locus was confirmed. Analysis of individual blastocysts derived from Cas9 RNP-electroporated zygotes revealed indels in all embryos (*SI Appendix*, Fig. S2A). Moreover, 86% of blastocysts displayed high editing levels, with 76–100% of alleles carrying indels, indicating that gRNA induces DSBs with exceptionally high efficiency (*SI Appendix*, Fig. S2A). Subsequently, a panel of ATM inhibitors in pig zygotes was screened by electroporating them with Cas9 RNP, followed by 24-h incubation with AAV donors and inhibitors. Although the variability was considerable, all ATM inhibitors showed a general trend toward improving knock-in efficiency in blastocysts (*SI Appendix*, Fig. S2B-H). However, with the exception of AZ32 and KU55933, all ATM inhibitors reduced blastocyst development at concentrations that appeared effective for enhancing knock-in. In contrast, treatment with AZ32 or KU55933 at 1 μM increased knock-in efficiency without a detectable decrease in blastocyst development. Because AZ32 showed a greater increase in knock-in efficiency than KU55933, it was selected for subsequent optimization.

To determine the optimal AZ32 concentrations, multiple concentrations were compared and their effects on blastocyst development and knock-in efficiency were assessed. AZ32 did not reduce blastocyst development significantly at any of the concentrations tested. In contrast, knock-in efficiency was improved significantly at 1–10 μM compared with in the untreated controls, with no appreciable differences among the doses (Fig. 2A and B). Based on the results, 1 μM was selected as the working concentration for subsequent experiments. Subsequently, whether AZ32 would still enhance knock-in under conditions that mimic multiplex knock-in, where the total AAV load must be shared among several donor vectors, and the dose of each donor is accordingly reduced, was examined. To evaluate that, electroporated zygotes were cultured with 1 μM AZ32 while reducing the AAV donor concentration progressively from 3×10^9^ to 0.25×10^9^ vg/mL (6×10^6^ to 0.5×10^6^ vg/zygote). AZ32 increased knock-in efficiency significantly at doses down to 0.5×10^9^ vg/mL and exhibited a similar trend even at 0.25×10^9^ vg/mL (*P* = 0.06), without any detectable effects of AZ32 on blastocyst development at any AAV dose tested (Fig. 2C). To investigate the underlying mechanism of the observed increase in knock-in efficiency, Simple Western analysis was performed 6 h after electroporation (Fig. 2D). AAV transduction resulted in marked activation of ATM, whereas the p53–caspase pathway showed no appreciable increase, in contrast to previous observations of the authors in pluripotent stem cells and hematopoietic stem cells (Natsagdorj et al., *Commun Biol* in Revision) (34). Instead, AAV transduction clearly increased DNA-PKcs phosphorylation at Thr2609, indicating predominant activation of the c-NHEJ pathway. In addition, treatment with AZ32 attenuated AAV-induced ATM and DNA-PKcs phosphorylation, indicating that ATM inhibition suppressed the AAV-induced shift toward c-NHEJ.

**Figure 2.**
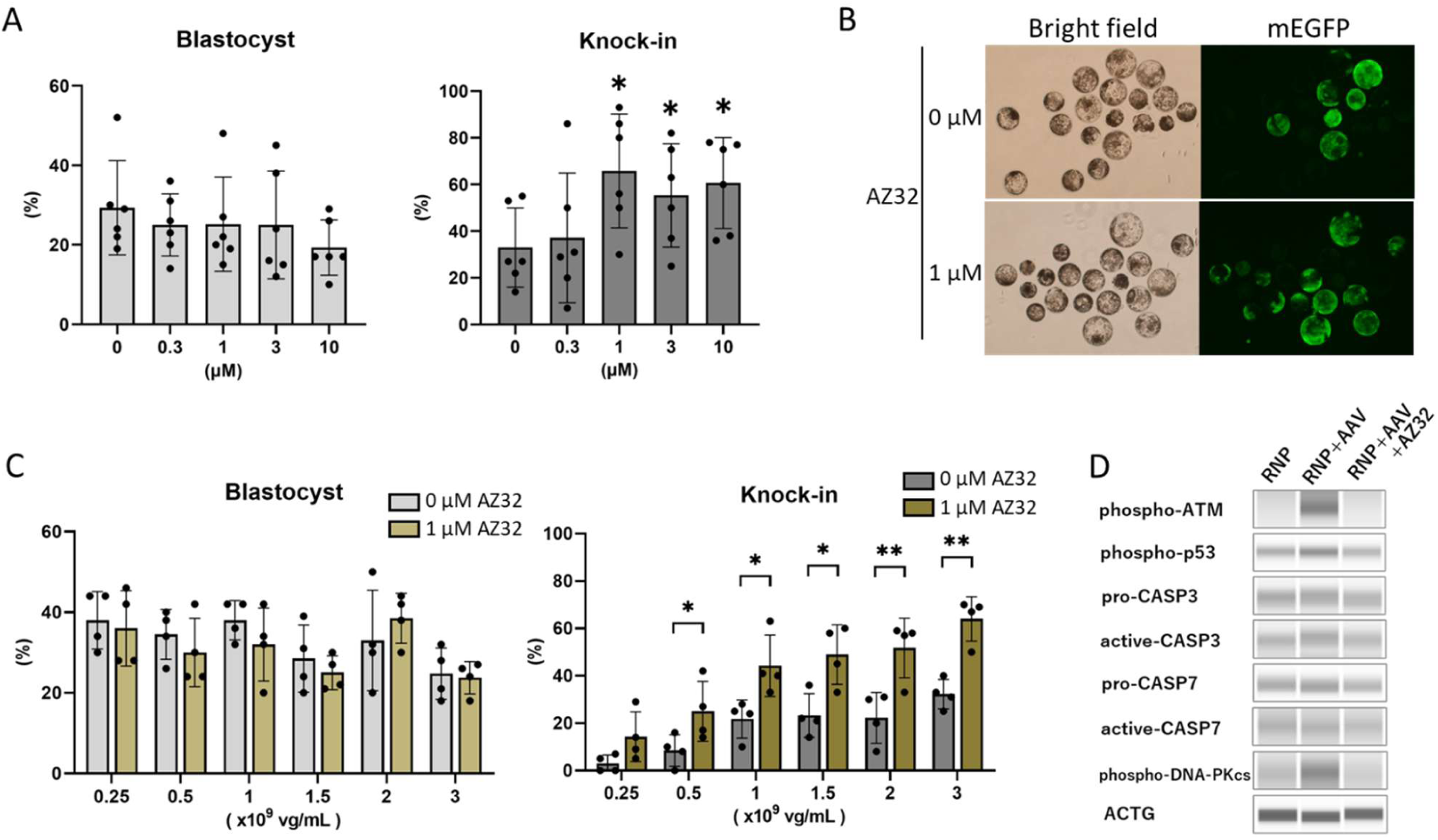
Improvement of adeno-associated virus (AAV)-mediated knock-in efficiency in pig zygotes by ataxia-telangiectasia mutated (ATM) inhibition. (*A*) Effect of AZ32 concentration on blastocyst development and on ACTB–mEGFP knock-in efficiency assessed at the blastocyst stage. Zygotes were electroporated with Cas9 RNP and transduced with an AAV donor (3×10^9^ vg/mL; 6×10^6^ vg/zygote). Bars show mean ± SD, and dots represent independent replicates. Data were analyzed by repeated measures one-way Analysis of Variance with the Geisser–Greenhouse correction. For follow-up analysis, Dunnett’s multiple comparisons test was used to compare each treatment with the control (n = 5 biological replicates). \**P* < 0.05. (*B*) Representative bright-field and mEGFP fluorescence images of blastocysts treated with 0 or 1 μM AZ32. (*C*) Effect of AAV donor dose on blastocyst development (left) and knock-in efficiency (right) in the presence or absence of 1 μM AZ32. Electroporated zygotes were cultured for 24 h with the indicated AAV doses (0.25–3 × 10^9^ vg/mL; 0.5–6 × 10^6^ vg/zygote) with or without 1 μM AZ32. Statistical significance between AZ32-treated and untreated samples at each AAV dose was evaluated by paired t-tests with Holm–Šidák correction for multiple comparisons (n = 4 biological replicates). \*\**P* < 0.01; \**P* < 0.05. (*D*) Simple Western analysis of DNA damage–response signaling 6 h after RNP electroporation. Lysates from zygotes treated with Cas9 RNP alone, RNP plus AAV (3 × 10^9^ vg/mL), or RNP plus AAV plus AZ32 (1 μM) were probed for phosphorylated ATM, phosphorylated p53, caspase-3 (CASP3), caspase-7 (CASP7), phosphorylated DNA-PKcs, and ACTG as a loading control.

### Efficient Generation of Knock-in Fetuses from ATM Inhibitor-Treated Embryos

Whether AZ32 treatment enabled efficient generation of knock-in fetuses *in vivo* was evaluated by transferring treated embryos into surrogate sows. A total of 12 E32 fetuses with normal development were retrieved, of which nine (75%) exhibited clear mEGFP fluorescence (Table 1, Fig. 3A), closely matching the knock-in efficiencies observed *in vitro* (Fig. 2A). All mEGFP-positive fetuses showed 5’ and 3’ junction PCR bands of the expected sizes for knock-in (Fig. 3B). Subsequently, porcine embryonic fibroblasts (PEFs) derived from each fetus were evaluated using flow cytometry. Consistent with the whole-mount fluorescence and PCR results, mEGFP-positive cell populations were present in all nine fluorescent fetuses (Fig. 3C). Notably, six fetuses consisted entirely of mEGFP-positive cells, and three of the fetuses displayed a population with higher fluorescence intensity, suggesting the presence of homozygous knock-in cells. Overall, fetuses derived from AZ32-treated zygotes exhibited high knock-in efficiencies without any clear signs of treatment-related abnormalities.

**Figure 3.**
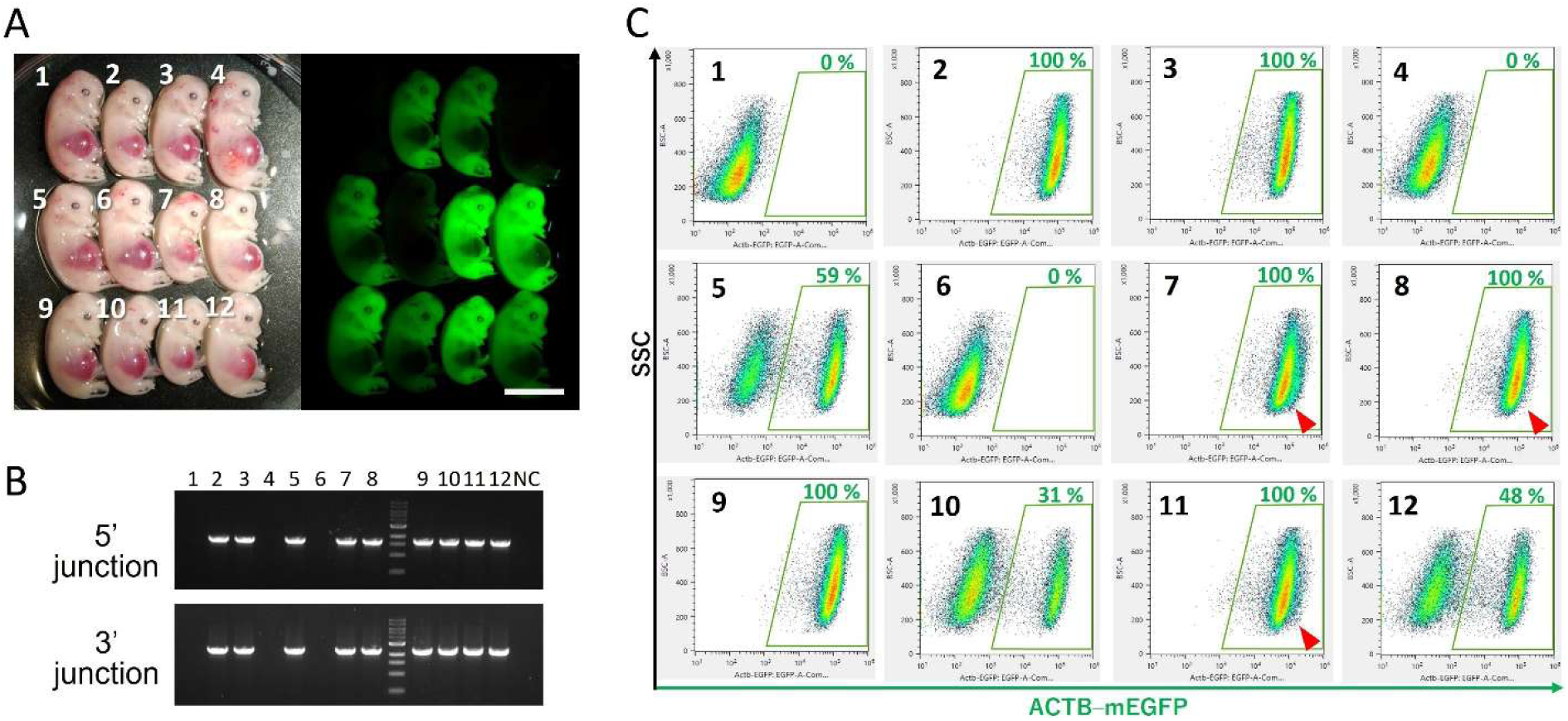
Efficient generation of ACTB–mEGFP knock-in fetuses following AZ32 treatment. (*A*) Whole-mount bright-field and mEGFP fluorescence images of E32 fetuses recovered from two separate surrogate sows after transfer of embryos that had been electroporated with Cas9 RNP and subsequently cultured with adeno-associated virus donor (3×10^9^ vg/mL) and 1 µM AZ32. Numbers correspond to individual fetuses. Scale bar, 2 cm. (*B*) Junction PCR analysis of the 5’ and 3’ knock-in junctions in genomic DNA from each fetus and a negative control (NC). (*C*) Flow cytometric analysis of pig embryonic fibroblasts derived from each fetus. Gates indicate mEGFP-positive cells, and percentages indicate the fraction of mEGFP-positive cells. Red arrowheads highlight a higher-fluorescence population suggestive of homozygous knock-in cells.

**Table 1.** ACTB–mEGFP knock-in efficiency in pig fetuses derived from AZ32-treated zygotes.

| No. of sows |  | No. (%) of fetuses |  |  | No. (%) of mEGFP-positive normal fetuses |
| --- | --- | --- | --- | --- | --- |
| Transferred | Pregnant | Total | Normal | Delayed |  |
| 2 | 2 | 17 | 12 (71) | 5 (29) | 9 (75) |

### Improvement of Simultaneous Double Knock-in at *ACTB* and *UBB* loci in Pig Zygotes by AZ32

To examine whether AZ32 also enhances multiple knock-in in pig zygotes, a second knock-in reporter system targeting the pig *UBB* locus was established. In the system, a *UBB*-targeting gRNA and an AAV donor that inserted an mScarlet-I3 cassette linked via a linker–2A sequence at the C-terminus of the endogenous *UBB* coding sequence was designed (Fig. 4A). The reporter allows independent visualization of knock-in at the *UBB* locus, thereby enabling identification of double knock-in blastocysts expressing both fluorescent proteins. Before applying the reporter in zygotes, the UBB–mScarlet-I3 system was validated in immortalized porcine MSCs. Similar to in the ACTB–mEGFP reporter, *UBB*-targeting Cas9 RNP alone or the UBB–mScarlet-I3 AAV donor alone produced no mScarlet-I3-positive cells, whereas their combination yielded a distinct mScarlet-I3–positive population (*SI Appendix*, Fig. S3A). Junction PCR of the sorted mScarlet-I3-positive and - negative fractions further confirmed that correct 5’ and 3’ knock-in junctions were present only in the positive fraction (*SI Appendix*, Fig. S3B). Based on the results, the reporter was tested in pig zygotes and whether *UBB*-targeting Cas9 RNP efficiently induced DSBs was assessed. Analysis of individual blastocysts derived from zygotes electroporated with the *UBB*-targeting Cas9 RNP showed that indels were detected in all embryos, and most blastocysts fell into the highest editing category, with 76–100% of alleles carrying indels (Fig. 4B). Thus, Cas9 RNP targeting *UBB* induces DSBs with high efficiency, similar to that observed previously at the *ACTB* locus. Next, whether AZ32 improves AAV-mediated knock-in at the *UBB* locus was tested. Zygotes were electroporated with *UBB*-targeting Cas9 RNP and cultured with the UBB–mScarlet-I3 AAV donor with or without 1 μM AZ32. Similar to in the ACTB–mEGFP system, AZ32 did not reduce blastocyst development but increased the proportion of UBB–mScarlet-I3-positive blastocysts significantly (Fig. 4C).

**Figure 4.**
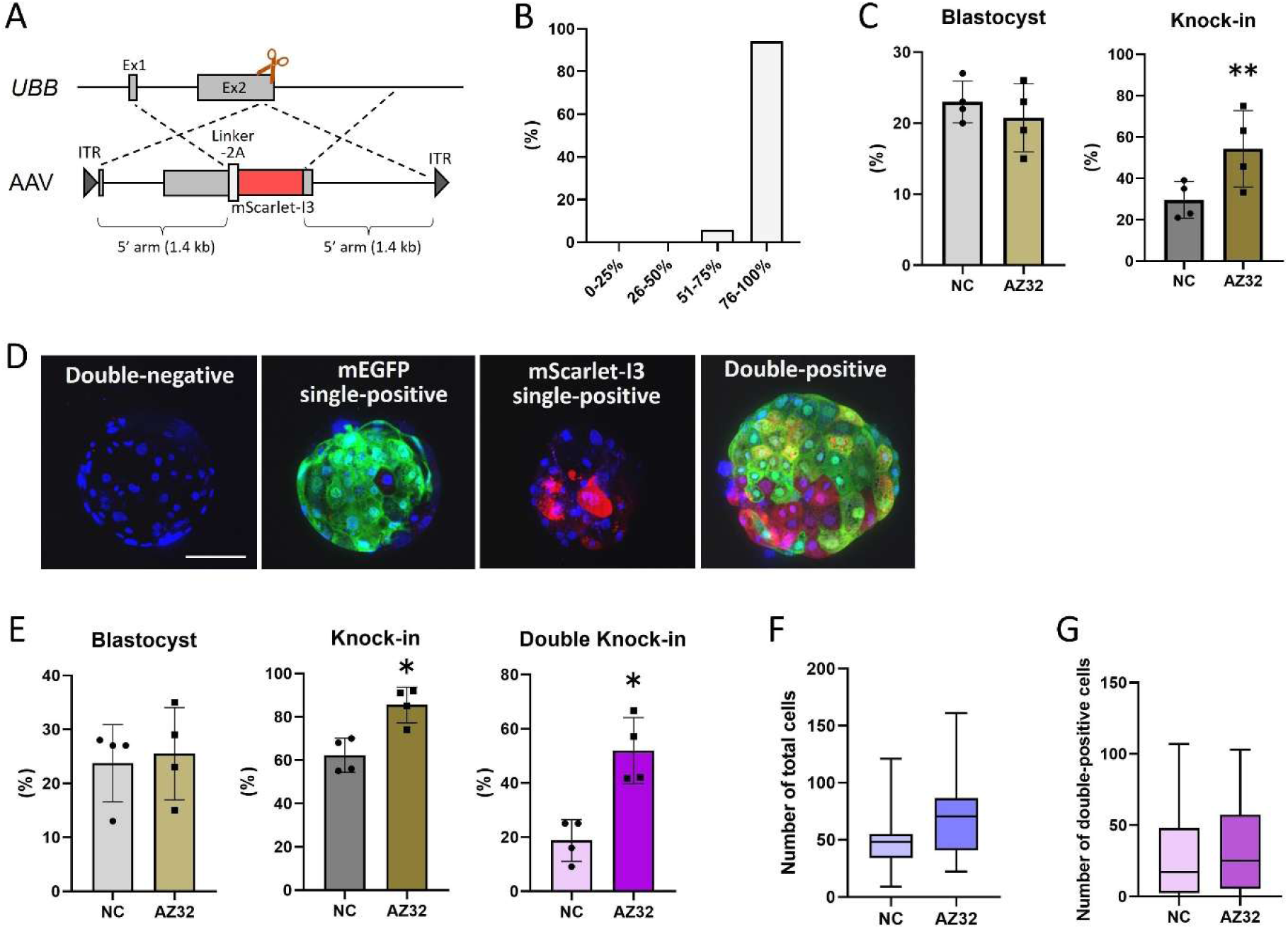
Improvement of UBB–mScarlet-I3 knock-in and simultaneous double knock-in at *ACTB* and *UBB* loci in pig zygotes by AZ32. (*A*) Schematic of the UBB–mScarlet-I3 knock-in strategy. An adeno-associated virus (AAV) donor carrying an mScarlet-I3 cassette linked via a linker–2A sequence and flanked by 1.4-kb 5’ and 3’ homology arms was designed to insert mScarlet-I3 in frame at the C terminus of the pig UBB coding sequence. The orange scissors indicate the Cas9 cleavage site in exon 2. ITR, inverted terminal repeat. (*B*) Distribution of indel frequencies at the *UBB* locus in individual blastocysts derived from zygotes electroporated with *UBB*-targeting Cas9 RNP. Genomic DNA from 17 blastocysts was isolated, the *UBB* target site was PCR amplified, Sanger sequenced, and indel frequencies were estimated by TIDE. Blastocysts were grouped into four classes according to the percentage of alleles carrying indels (0–25, 26–50, 51–75, and 76–100%). (*C*) Effect of AZ32 on blastocyst development (left) and UBB–mScarlet-I3 knock-in efficiency (right). Zygotes were electroporated with *UBB*-targeting Cas9 RNP and cultured for 24 h with UBB–mScarlet-I3 AAV donor (3 × 10^9^ vg/mL; 6 × 10^6^ vg/zygote) in the presence or absence of 1 μM AZ32, then cultured in inhibitor-free medium for an additional 6 d. Bars show mean ± SD, and dots represent biological replicates (n = 4). Statistical significance between NC and AZ32 conditions was evaluated by paired t tests. \*\**P* < 0.01. (*D*) Representative fluorescence images of blastocysts classified as double negative, ACTB–mEGFP single-positive, UBB–mScarlet-I3 single-positive, or double-positive for both reporters under double knock-in conditions. Blue, nuclear DNA; green, ACTB–mEGFP; red, UBB–mScarlet-I3. Scale bar, 100 μm. (*E*) Effect of AZ32 on simultaneous double knock-in at the *ACTB* and *UBB* loci. Zygotes were electroporated with Cas9 RNPs targeting both *ACTB* and *UBB* (1 μM each) and cultured with both AAV donors (1.5 × 10^9^ vg/mL each; total 3 × 10^9^ vg/mL) in the presence or absence of 1 μM AZ32. The panels show blastocyst development (left), the percentage of blastocysts with knock-in at either locus (middle), and the percentage of double knock-in blastocysts (right). Bars show mean ± SD, and dots represent biological replicates (n = 4). Statistical significance between NC and AZ32 conditions was evaluated by paired t tests. \**P* < 0.05. (F and G) Quantification of total cell numbers (F) and double-positive cell numbers (G) in double-positive blastocysts obtained under double knock-in conditions with or without AZ32. Box plots indicate median, interquartile range, and minimum and maximum values. No significant differences were detected between NC and AZ32 groups by unpaired t-tests (NC, n = 17; AZ32, n = 32).

Subsequently, simultaneous double knock-in at the *ACTB* and *UBB* loci was examined. To keep the total amounts of Cas9 RNPs and AAV donors equivalent to those used for single knock-in, a half of the single-locus amounts was used for each locus (1 μM for each Cas9 RNP and 1.5×10^9^ vg/mL for each AAV donor). Under the conditions, AZ32, once more, had no detectable effect on blastocyst development, but it increased the percentage of blastocysts in which knock-in occurred at either locus and increased the percentage of blastocysts that were double-positive for ACTB–mEGFP and UBB–mScarlet-I3 (Fig. 4D and E). Total numbers of cells in the resulting double-positive blastocysts were quantified and the number of double knock-in cells within each blastocyst was assessed. The total cell numbers were comparable between the AZ32-treated and untreated groups (Fig. 4F). The number of double knock-in cells per double-positive blastocyst was similar between the two groups (Fig. 4G). Overall, the results indicate that AZ32 enhances the yield of double knock-in blastocysts without compromising their quality.

### Generation of Double Knock-in Fetuses from AZ32-Treated Zygotes, Including an F0 Fetus Composed Almost Entirely of Double Knock-in Cells

To determine whether AZ32-treated zygotes could give rise to double knock-in fetuses in the F0 generation, embryos treated under double knock-in conditions were transferred into three surrogate sows. Embryo transfer yielded nine fetuses with normal development at E32–34. Whole-mount fluorescence imaging showed reporter expression in six fetuses: five expressed ACTB–mEGFP, four expressed UBB–mScarlet-I3, and three expressed both reporters (Table 2, Fig. 5A). Junction PCR results supported the observations, with ACTB–mEGFP 5’ and 3’ junctions detected in five fetuses and UBB–mScarlet-I3 junctions detected in four (Fig. 5B). In total, six of the nine fetuses carried knock-in alleles at either locus, and three fetuses carried alleles at both loci. To corroborate the findings at the cellular level, PEFs from each fetus were examined using flow cytometry (Fig. 5C). The corresponding fluorescent cell population was also detected by flow cytometry in every fetus identified as knock-in positive by imaging and PCR. Notably, one fetus (#1) consisted almost entirely of double-positive PEFs, indicating that the fetus was composed predominantly of double knock-in cells. The results demonstrate that AZ32-treated zygotes can efficiently generate single and double knock-in fetuses and suggest that producing F0 animals composed almost entirely of double knock-in cells is now feasible.

**Figure 5.**
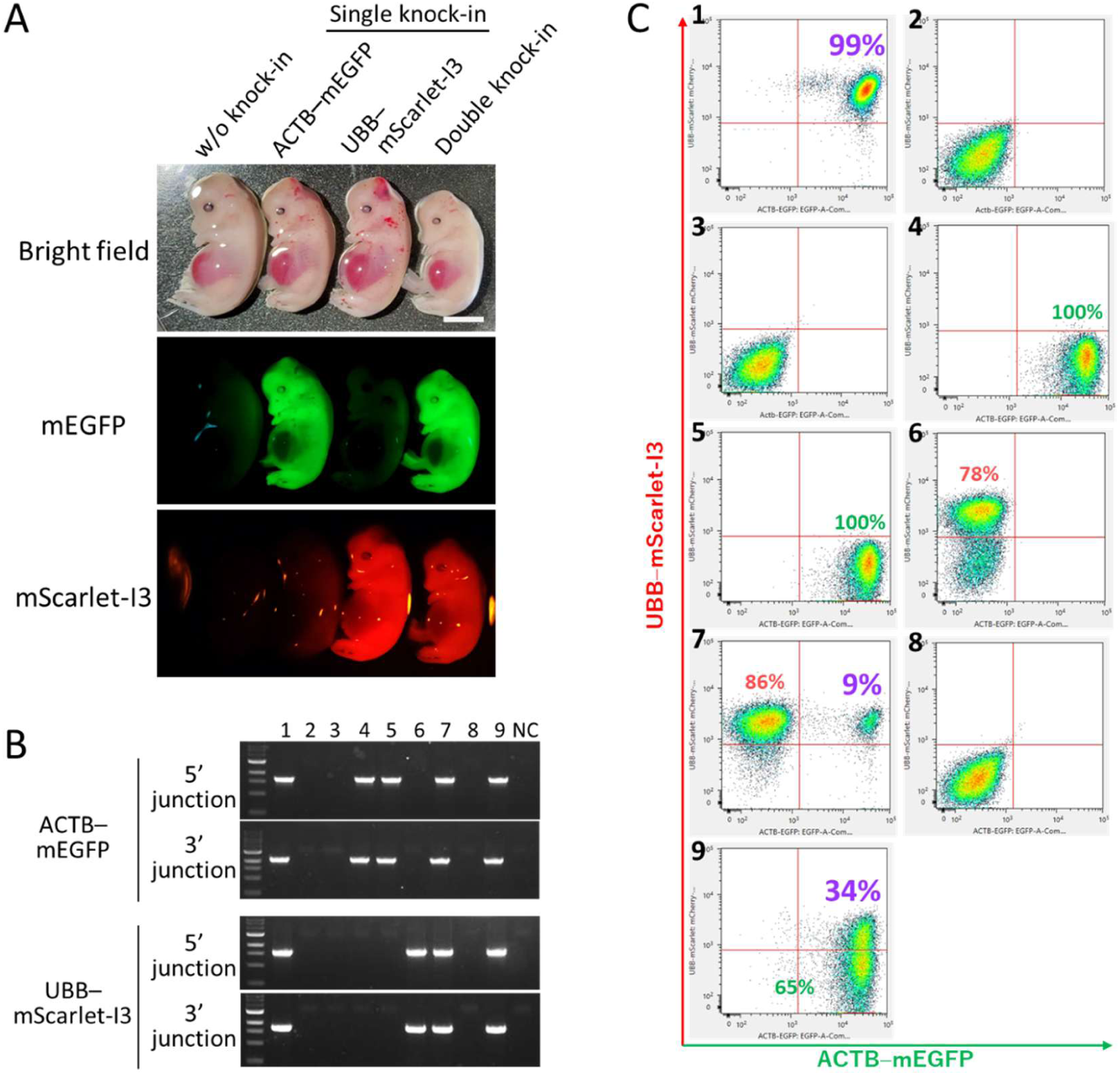
Efficient generation of double knock-in fetuses from AZ32-treated zygotes. (*A*) Representative bright-field, ACTB–mEGFP, and UBB–mScarlet-I3 fluorescence images of E32-34 fetuses obtained after transfer of embryos that had been cultured under double knock-in conditions in the presence of 1 μM AZ32. Scale bar, 1 cm. (*B*) Junction PCR analysis of ACTB–mEGFP and UBB–mScarlet-I3 knock-in alleles in fetuses #1-#9 and a negative control (NC). (*C*) Flow cytometry analysis of pig embryonic fibroblasts derived from each fetus. Numbers 1–9 correspond to the fetuses shown in panel B, and percentages indicate the fractions of cells in each fluorescence gate (single-positive, and double-positive). Fetus #1 consists almost entirely of double-positive cells, indicating that this F0 animal is composed predominantly of double knock-in cells.

**Table 2.** ACTB–mEGFP and UBB–mScarlet-I3 double knock-in efficiency in pig fetuses derived from AZ32-treated zygotes.

| No. of sows |  | No. (%) of fetuses |  |  | No. (%) of normal fetuses* |  |  |  |
| --- | --- | --- | --- | --- | --- | --- | --- | --- |
| Transferred | Pregnant | Total | Normal | Delayed | Either-reporter-positive | mEGFP-positive | mScarlet-I3-positive | Double-positive |
| 3 | 3 | 16 | 9 (56) | 7 (44) | 6 (67) | 5 (56) | 4 (44) | 3 (33) |

\* Percentages are based on normally developed fetuses.

## Discussion

A key advantage of zygote-based genome editing is its ability to generate genome-engineered pigs directly without relying on SCNT, which is often associated with epigenetic abnormalities. In the present study, the authors demonstrate that AAV donor vectors support efficient long-fragment knock-in in pig zygotes, and that suppressing AAV-induced ATM overactivation with AZ32 further enhances knock-in efficiency without impairing embryo development and blastocyst quality. Such advances enabled efficient long-fragment knock-in in pigs for the first time without the need for selection (29). Furthermore, AZ32 increased the efficiency of simultaneous double knock-in and enabled the generation of double knock-in fetuses, including an F0 fetus composed almost entirely of double knock-in cells. The findings demonstrate that combining AAV donors with suppression of AAV-induced ATM overactivation is a practical approach for achieving multiple knock-in alleles in a single generation, which in turn, enables one-step generation of conditional knockout models for studying tissue-specific gene function, organ-deficient hosts for producing human organs through interspecies organogenesis, and other genetically complex configurations required for advanced applications in pigs. However, combining AAV donors with ATM inhibition yielded mosaic blastocysts and fetuses containing wild-type and single knock-in cells (Fig. 4D and 5B). Reducing such mosaic contributions, in addition to obtaining full-term offspring and assessing their normal development, are vital for strengthening the broader applicability of the approach. Nevertheless, the strategy remains highly valuable for generating genetically engineered pigs, in contrast to rodent models, where multiple single-locus knock-in lines can be assembled through intercrossing, because establishing and intercrossing such lines in pigs requires substantial time and cost resources.

AAV vectors exhibit low immunogenicity (36), deliver DNA of up to approximately 4.7 kb into mammalian cells with high efficiency (37, 38), and rarely undergo random genomic integration (39–41). Consequently, their use as donors for long-fragment knock-in has expanded rapidly across diverse cell types, including rodent zygotes (30, 32, 42–44). However, their effectiveness as donor vectors in large-animal zygotes has not yet been examined. Our results demonstrated that AAV donors enabled successful fluorescent protein knock-in at both *ACTB* and *UBB* loci in pig zygotes and that embryos carrying the knock-in alleles progressed through early development. The proportion of blastocysts containing knock-in cells was approximately 30% (Fig. 2), which was insufficient for reliable multi-locus knock-in.

ATM is a central sensor kinase in the DNA damage response and plays a well-established role in DSB repair by phosphorylating key recombination factors, such as CtIP, BRCA1, and RAD51 (45–47). We have recently showed that AAV and other linear DNA donors activate ATM strongly in pluripotent stem cells, which in turn triggers two major downstream responses: activation of the p53–caspase apoptosis pathway and stimulation of DNA-PKcs-dependent c-NHEJ, a principal repair pathway that competes with knock-in (Natsagdorj et al., *Commun Biol*, in Revision). Additionally, pharmacological inhibition of ATM suppresses such responses and increases knock-in efficiency across various cell types (Natsagdorj et al., *Commun Biol*, in Revision) (34). Consistent with the findings, ATM inhibitors increased AAV-mediated knock-in in porcine zygotes, including double knock-in (Fig. 2 and 4). However, mechanistic analysis in zygotes revealed a distinct profile. AAV transduction caused strong phosphorylation of ATM and DNA-PKcs at the T2609 cluster, in agreement with previous reports that the site is phosphorylated in an ATM-dependent manner (48), whereas activation of the p53–caspase pathway was minimal (Fig. 2D). The findings suggest that apoptosis is not the major barrier in pig zygotes, and that ATM primarily limits AAV-mediated knock-in by promoting c-NHEJ through DNA-PKcs. The difference from our previous studies using mouse pluripotent stem cells may reflect the fact that the actual amount of AAV donor transduced into zygotes is insufficient to trigger robust apoptosis or that the ATM–p53 axis is inherently less responsive in zygotes. Notably, the observation that only a subset of ATM inhibitors, such as AZ32 and KU55933, enhanced knock-in without reducing blastocyst development, may reflect differences in compound-specific properties, such as the persistence of inhibitory activity after reagent removal or distinct off-target effects.

One of the key technical elements of the present study is the development of robust reporter systems for quantifying knock-in in pig blastocysts. Since *in vivo* experiments in pigs are costly and cannot be performed at high throughput, it is essential to establish *in vitro* assays that provide sufficient signal-to-noise ratios for clear detection of knock-in at the blastocyst stage. We first selected the ACTB–mEGFP reporter based on our previous work in mouse cells, which demonstrated that fusing mEGFP to the C-terminus of ACTB yields bright and stable fluorescence (Natsagdorj et al., *Commun Biol*, in Revision) (34). The design was applied to pigs and confirmed robust mEGFP expression in porcine MSCs, blastocysts, and fetuses (Fig. 1-3). Second, *UBB*, which is also a reference gene, was targeted. In pigs, UBB is expressed highly, with low variability across multiple tissues, and has been validated as a reliable normalization marker for expression studies (49, 50). Additionally, our reanalysis of the publicly available pig embryo RNA-seq dataset (GSE139512) from Kong et al. (51) confirmed that UBB is expressed broadly and exhibits stable expression throughout preimplantation development, including the blastocyst stage. For the UBB knock-in reporter, mScarlet-I3 (52), which is a recently reported member of the mScarlet fluorescent protein family (53), was used. Although mCherry has long been favored owing to its low cytotoxicity, its brightness, defined as the product of the extinction coefficient and quantum yield, is approximately a half that of mEGFP. mScarlet-I3 provides more than four–fold higher brightness than mCherry, while exhibiting lower cytotoxicity than both mCherry and mEGFP (52), making it well-suited for conditions where reporter expression is high and cytotoxicity can become limiting. The ACTB–mEGFP and UBB–mScarlet-I3 combination yields spectrally separable high-intensity signals that allow robust discrimination of double-negative, single-positive, and double-positive blastocysts and fetuses (Fig. 4D, 5 B, and C). These reporters provide practical tools for improving multiplex knock-in protocols in pigs.

Pig cell lines generated in the present study by introducing the SV40 large T antigen share the same genetic background as the pigs used for gamete collection. The cells displayed surface marker profiles characteristic of MSCs (*SI Appendix* Fig. S1C) (35, 54). Because these cells are readily transduced by AAV6, the same serotype used in the zygote experiments, they provide a suitable platform for pre-evaluating knock-in reporter systems that can be robustly and quantitatively assessed by fluorescence-based cell sorting (Fig. 1B, C and *SI Appendix*, Fig. S3). Additionally, as experiments using zygotes exhibit substantial biological variability and cannot handle many experimental groups, MSC-based assays serve as a scalable platform for prioritizing compounds prior to zygote experiments. In such a cellular context, multiple ATM inhibitors produced dose-dependent increases in AAV-mediated ACTB–mEGFP knock-in, whereas CGK733 failed to enhance knock-in and instead reduced knock-in efficiency significantly. The finding is consistent with previous reports questioning whether CGK733 acts as a selective ATM inhibitor (55, 56). Notably, the optimum concentrations for most ATM inhibitors in zygotes were approximately one-tenth to one-hundredth of the highest concentrations tested in MSCs without causing complete cell loss, indicating that MSC-based assays can be used to approximate effective potency ranges before zygote experiments.

In summary, combining AAV donor vectors with transient inhibition of ATM by AZ32 enabled efficient long-fragment knock-in at single and multiple loci in porcine zygotes. The approach yielded double knock-in fetuses, including an F0 fetus composed almost entirely of double knock-in cells. The findings provide a practical route for introducing multiple targeted alleles into a single generation of pigs.

## Materials and methods

### Animals

All animal care and experimental procedures were performed in accordance with the Animal Experiment Guidelines of Jichi Medical University, Tochigi, Japan, and they were approved by the Animal Care Committee of Jichi Medical University (23037–04). The health of all pigs was monitored daily during feeding by animal husbandry staff under the supervision of attending veterinarians. Euthanasia was performed under deep sevoflurane anesthesia by intravenous injection of a potassium chloride solution (3 mmol/kg), in compliance with the American Veterinary Medical Association Guidelines for the Euthanasia of Animals.

### Establishment of Immortalized Pig MSCs

Bone marrow–derived primary adherent cells were obtained by culturing bone marrow collected from the humerus of domestic LWD (Landrace × Large White × Duroc) pigs. The bone marrow was flushed with ACD-A solution (JMS, Hiroshima, Japan), treated with ACK lysis buffer to remove red blood cells, filtered through a 70-µm strainer, and plated in MSC medium. MSC medium was composed of Minimum Essential Medium alpha (Gibco, FMD, USA) supplemented with 20% fetal bovine serum (FBS, Gibco) and 1× penicillin–streptomycin (Thermo Fisher Scientific, MA, USA). Primary adherent cells at early passage were electroporated with pEf1a-Large T antigen-IRES-Puro (Addgene #18922). Electroporation was performed using a NEPA21 electroporator (Nepagene, Chiba, Japan) and 2-mm electroporation cuvettes, with four poring pulses at 125 V for 2 ms, followed by five transfer pulses at 20 V for 50 ms. After electroporation, cells were cultured for 24 h and then selected with 1 µg/mL puromycin for 7 d. Puromycin-resistant cells were maintained in the MSC medium without puromycin. To determine whether immortalized cells exhibited MSC-like characteristics, surface marker profiling was performed using flow cytometry. The cells were detached using the Cell Dissociation Solution (Sigma-Aldrich, MO, USA) and stained with the antibodies listed in *SI Appendix* (Table S1). Dead cells were stained with DAPI (Dojindo, Kumamoto, Japan) immediately before analysis, and only DAPI-negative cells were subjected to flow cytometry using an SH800 cell sorter (Sony Biotechnology, CA, USA).

### Genome-Editing in Pig MSCs

Genome editing of porcine MSCs was performed by electroporation with Cas9 RNP followed by transduction with AAV donor vectors. To prepare the Cas9 RNP, the target-specific crRNA for either the pig *ACTB* locus (AGGTTTTGTCAAGAAAAAGG; IDT, IA, USA) or the *UBB* locus (GCATTCATAATGCCCACTGA; IDT) and the tracrRNA (IDT) were mixed, incubated for 5 min at 95°C, and cooled to room temperature to allow duplex formation. The duplex was then combined with Cas9 protein (IDT) at a final concentration of 18.6 µM and incubated for an additional 20 min at room temperature (57). Pig MSCs were electroporated with Cas9 RNP at a final RNP concentration of 0.744 µM using the same electroporation settings described above for cell immortalization. Immediately after electroporation, crude-grade AAV6 donor vectors (VectorBuilder) were added at 1×10^5^ vg/cell, and the cells were cultured for 24 h in MSC medium without FBS to avoid the inhibition of AAV transduction by FBS (58). When ATM inhibitors were tested, they were added during the same 24-h incubation. After 24 h, the medium was replaced with standard MSC medium, and the cells were cultured for an additional 48 h. Flow cytometry analysis was performed 72 h after electroporation using an SH800 cell sorter (Sony Biotechnology). For genotyping, genomic DNA was extracted using the DNeasy Blood and Tissue Kit (Qiagen, MD, USA), and the target loci were amplified by PCR using the primer sets listed in *SI Appendix*, Table S2.

### Genome-Editing in Pig Zygotes

Pig ovaries from prepubertal LWD gilts were obtained from a local slaughterhouse, and cumulus–oocyte complexes collected from these ovaries were cultured in maturation medium for 44 h, as described previously (28, 59). *In vitro* fertilization was then performed using pooled semen from three purebred Landrace boars obtained from the National Livestock Breeding Center, Ibaraki Station (Ibaraki, Japan) at a final concentration of 2×10^6^ sperm/mL. Eight hours after insemination, cumulus cells and residual sperm were removed from the presumptive zygotes by gentle pipetting. For electroporation, Cas9 RNP was prepared as described above. Zygotes were electroporated with Cas9 RNP at a final concentration of 2 μM using a CUY21EDIT II electroporator (BEX, Tokyo, Japan) with five 1 ms pulses at 25 V. Immediately after electroporation, zygotes were transferred into PZM-5 medium (Research Institute for the Functional Peptides Co., Yamagata, Japan) supplemented with 3 × 10^9^ vg/mL AAV6 donor (6 × 10^6^ vg/zygote) and the indicated ATM inhibitors, and cultured for 24 h at 39°C under 5% CO₂, 5% O₂, and 90% N₂. The zygotes were then washed and cultured for an additional 6 d in PZM-5. FBS was added at a final concentration of 10% on day 5 after electroporation. For indel quantification, genomic DNA was obtained from individual blastocysts by alkaline extraction, and amplicons generated using the corresponding primer sets (*SI Appendix*, Table S2) were subjected to Sanger sequencing. The resulting chromatograms were analyzed using TIDE (https://apps.datacurators.nl/tide/) to quantify indel frequencies. To assess the knock-in efficiencies and count cell numbers in double knock-in blastocysts, fluorescence imaging was performed using a BZ-X microscope (Keyence, Osaka, Japan).

### Simple Western Analysis in Pig Zygotes

Pig zygotes were collected 6 h after electroporation and 400 zygotes were processed per experimental group. Samples were washed in PBS(–) containing 0.1 mg/mL polyvinyl alcohol and lysed in 20 μL of RIPA buffer (Nacalai Tesque, Tokyo, Japan) supplemented with 1× Phosphatase Inhibitor Cocktail II and III (Sigma-Aldrich) and 1× Protease Inhibitor (Nacalai Tesque). An equal volume (20 μL) of 2× Laemmli buffer was added, followed by 4 μL of 1 M DTT, gentle vortexing, and heating at 95°C for 5 min. Lysates were analyzed using the JESS Simple Western System (ProteinSimple, CA, USA), according to the manufacturer’s instructions. Data were processed using the Compass for Simple Western software (Bio-Techne, MN, USA). Information on the primary antibodies is provided in the *SI Appendix* (Table S3).

### Embryo Transfer And Fetal Analysis

Recipient gilts were Landrace or Landrace × Large White animals aged 4.5–5 months that had not yet experienced their first estrus. To synchronize estrus, 1,000 IU of equine chorionic gonadotropin (ZENOAQ, Fukushima, Japan) was administered intramuscularly, followed 72 h later by intramuscular administration of 1,500 IU of human chorionic gonadotropin (Asuka Animal Health, Tokyo, Japan). Embryos were obtained from two batches of electroporated zygotes generated on two consecutive days. Only embryos that had reached the 4-cell stage in the first batch (approximately 46 h after electroporation) or the 2-cell stage in the second batch (approximately 22 h after electroporation) were selected and combined. A total of 250–300 embryos were surgically transferred into each recipient, with embryos divided equally between the two oviducts following a procedure described previously (60).

The fetuses were recovered from the uterus on embryonic days 32–34 (E32–E34). Whole-mount bright-field and fluorescence imaging was performed to assess ACTB–mEGFP and UBB–mScarlet-I3 expression. For genotyping, genomic DNA was isolated from fetal tissues using the DNeasy Blood and Tissue Kit (Qiagen), and the 5’ and 3’ knock-in junctions were amplified by PCR using the primer sets listed in *SI Appendix*, Table S2. PEFs were established from each fetus by plating the dissociated fetal tissues in DMEM supplemented with 10% FBS and 1× penicillin–streptomycin. After expansion, PEFs were analyzed using flow cytometry to quantify the fractions of ACTB–mEGFP-positive and UBB–mScarlet-I3-positive cells. Dead cells were stained with DAPI (Dojindo) immediately before analysis, and only DAPI-negative cells were subjected to flow cytometry using an SH800 cell sorter (Sony Biotechnology).

### Statistical analysis

Statistical analyses were performed using Prism version 8 (GraphPad Software CA, USA). Unless otherwise indicated, data are presented as mean ± standard deviation from independent biological experiments. Comparisons between two groups were performed using paired or unpaired t-tests, as appropriate, with the Holm–Šidák correction applied for multiple paired comparisons. For comparisons between more than two groups, repeated-measures one-way Analysis of Variance with Geisser-Greenhouse correction was used, followed by Dunnett’s multiple comparisons test versus the control group. Dose-response relationships for AAV-mediated knock-in in the presence of ATM inhibitors were analyzed using linear regression of log-transformed inhibitor concentrations against knock-in frequencies, and statistical significance was evaluated by testing whether the regression slope differed from zero. *P* values < 0.05 were considered statistically significant. The statistical tests used, exact n values, and details of the multiple comparison procedures are provided in the figure legends.

## Acknowledgments

We thank the animal husbandry staff of the Animal Resource Laboratory at the Center for the Development of Advanced Medical Technology, Jichi Medical University, for their expert care and management of the pigs used in this study. This work was supported in part by the Japan Agency for Medical Research and Development (AMED) under Grant Number JP22bm1223003 to A.H. and F.T.. In addition, Y.H. and H.H. received research funding from Sumitomo Pharma Co., Ltd. through a Collaborative Research Agreement between Jichi Medical University and Sumitomo Pharma Co., Ltd.

## Author Contributions

H.H. and A.H. designed the research; H.H., M.N., F.T., and A.H. conducted the experiments; H.H. analyzed the data and wrote the original draft; M.N., F.T., M.I., Y.H., and A.H. reviewed and edited the manuscript.

