## Supporting Information for "Ataxia-Telangiectasia Mutated Inhibition Enhances Adeno-Associated Virus-Mediated Knock-in Efficiency in Pig Zygotes"

Arata Honda, PhD;

Hiromasa Hara, PhD;

**Supplementary Figures**

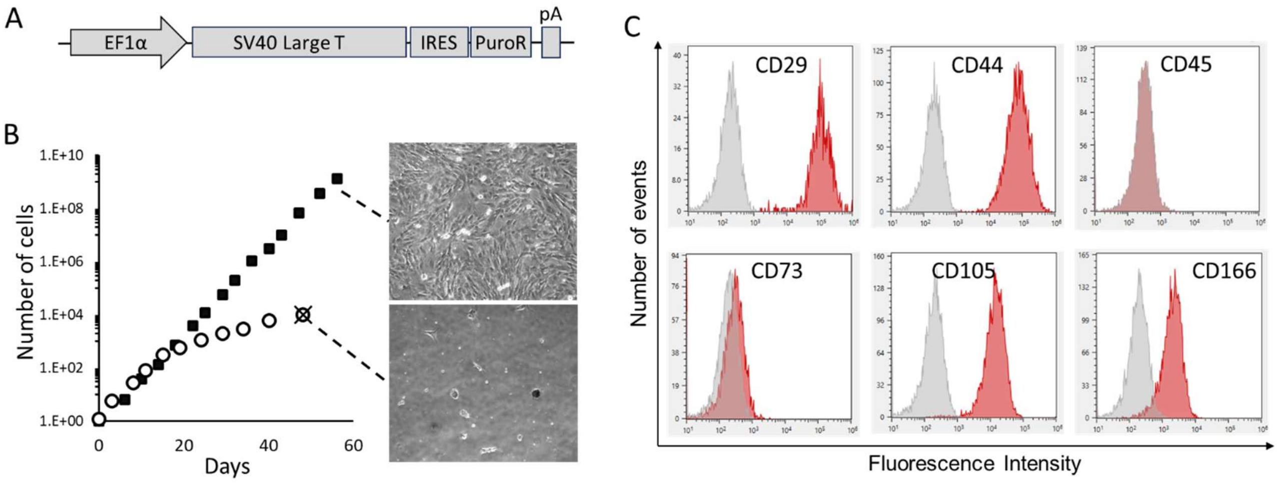


**Fig. S1.** Establishment and characterization of immortalized pig mesenchymal stromal cells (MSCs). (*A*) Schematic representation of the SV40 large T antigen expression cassette (pEF1α–SV40 large T–IRES–PuroR–pA) used to immortalize bone marrow-derived adherent cells. (*B*) Growth curves of primary (open circles) and immortalized (filled squares) pig bone marrow-derived cells. Representative phase-contrast images of proliferating immortalized cells (upper panel) and primary cells after growth arrest (lower panel) are shown on the right. (*C*) Flow cytometry histograms of surface marker expression in the immortalized cells. Red histograms indicate staining for MSC-associated markers (CD29, CD44, CD45, CD73, CD105, and CD166), and gray histograms indicate negative controls. For CD45, an isotype-matched control antibody was used as the negative control, whereas for all other markers, the negative controls were obtained by omitting the primary antibody.

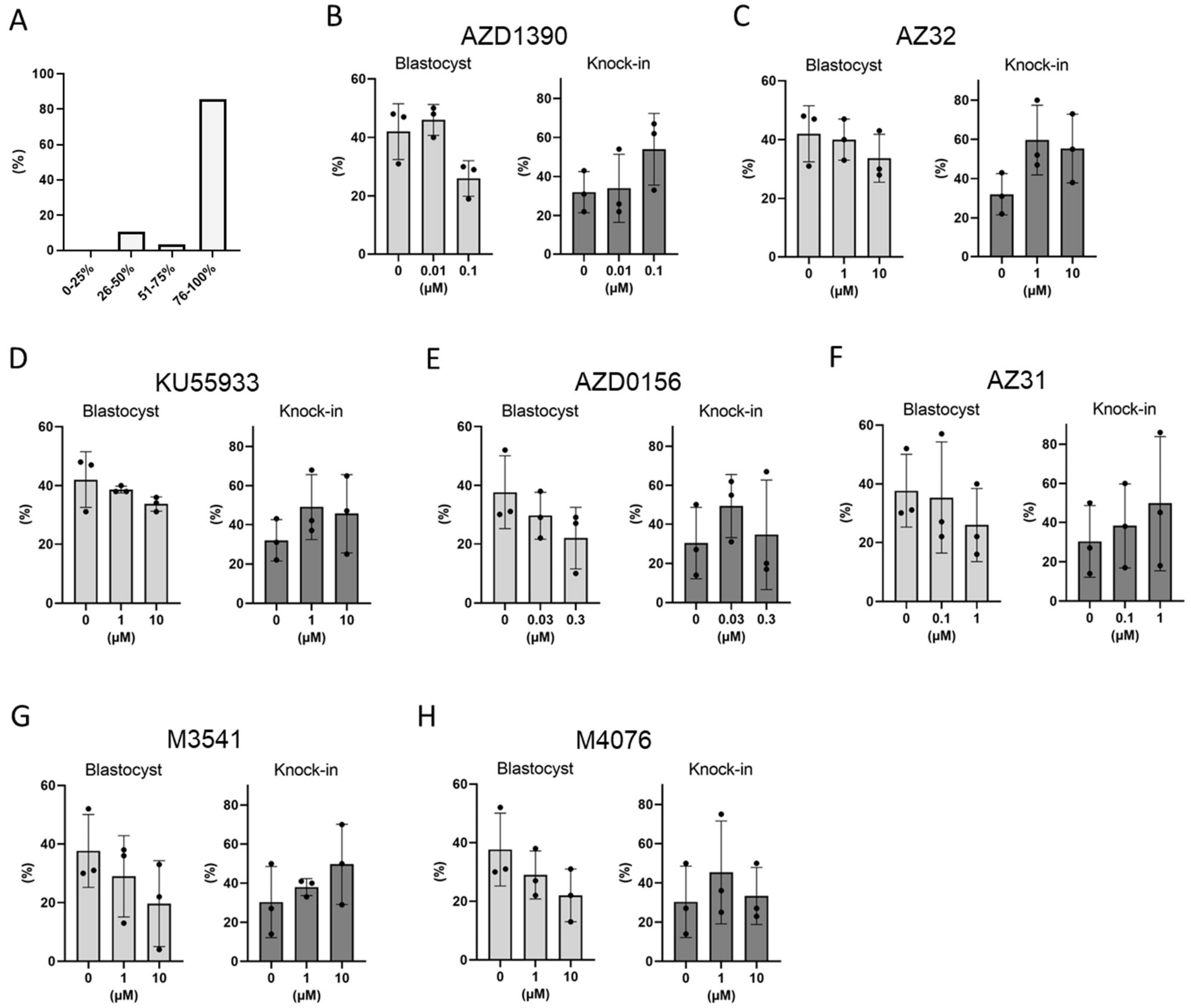


**Fig. S2.** Screening of ataxia-telangiectasia mutated (ATM) inhibitors in pig zygotes. (*A*) Distribution of indel frequencies at the *ACTB* locus in 28 individual blastocysts derived from zygotes electroporated with Cas9 RNP. For each blastocyst, genomic DNA was isolated, the *ACTB* target site was PCR amplified and Sanger sequenced, and the indel frequency was estimated by TIDE. Blastocysts were then grouped into four classes according to the percentage of alleles carrying indels (0–25, 26–50, 51–75, and 76–100%). (*B*–*H*) Effects of the indicated ATM inhibitors on blastocyst development and ACTB–mEGFP knock-in efficiency. Zygotes were electroporated with Cas9 RNP, then cultured for 24 h in the presence of AAV donor (3 × $10^9$ vg/mL) and the indicated concentrations of each inhibitor, followed by culture in inhibitor-free medium for an additional 6 d. Bars show mean ± SD. In panels B–D and in panels E–H, the 0 μM control data were obtained from the same experimental batches and were therefore shared across the inhibitors within each group (n = 3 biological replicates).

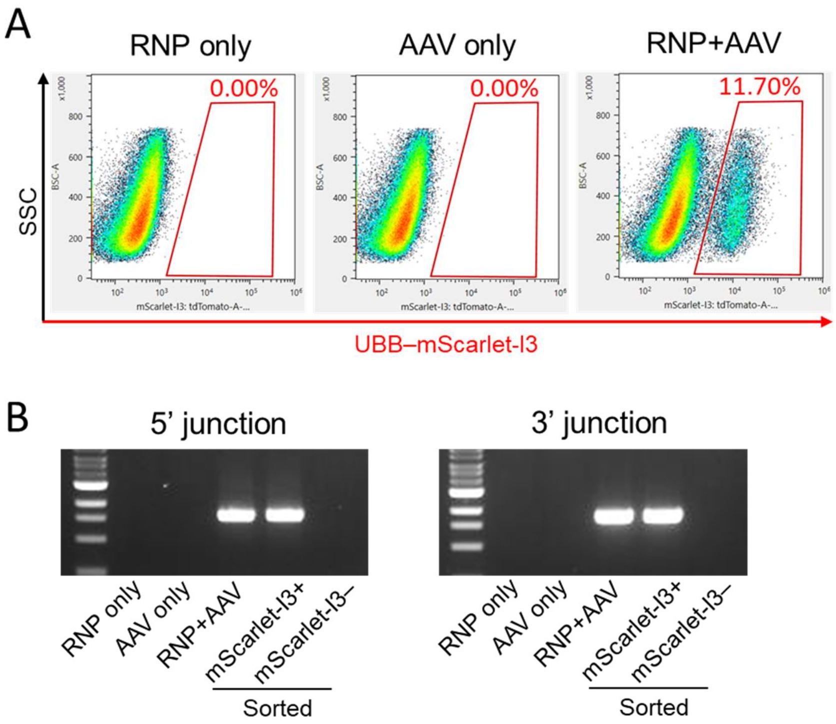


**Fig. S3.** Validation of the UBB–mScarlet-I3 knock-in reporter in immortalized pig mesenchymal stromal cells (MSCs). (*A*) Representative flow cytometry plots of immortalized pig MSCs 72 h after treatment with Cas9 RNP alone (RNP only), UBB–mScarlet-I3 AAV donor alone (AAV only), or both together (RNP + AAV). The red gate indicates UBB–mScarlet-I3–positive cells, and percentages denote the fraction of gated cells. (*B*) Junction PCR analysis of UBB–mScarlet-I3 knock-in alleles.

### Supplementary Tables

**Table S1.** Antibodies used for flow cytometry.

| No | Name of the antibody | Manufacturer | Cat# |
|---|---|---|---|
| 1 | Human/Porcine/Equine Integrin beta 1/CD29 MAb (Clone 419127) | R&D Systems | MAB17783-SP |
| 2 | CD44 antibody \| MAC329 | Bio-Rad | MCA1449GA |
| 3 | Mouse anti Pig CD45:Alexa Fluor® 647 | Bio-Rad | MCA1222A647 |
| 4 | Mouse IgG1 Negative Control:Alexa Fluor® 647 | Bio-Rad | MCA928A647 |
| 5 | Mouse/Porcine 5'-Nucleotidase/CD73 Affinity Purified PAb | R&D Systems | AF4488-SP |
| 6 | CD105 Monoclonal Antibody (MEM-229) | Invitrogen | MA1-19408 |
| 7 | CD166 Polyclonal Antibody | Bioss | bs-1251R |
| 8 | F(ab')2-Goat anti-Rat IgG (H+L) Cross-Adsorbed Secondary Antibody, PE | Invitrogen | A10544 |
| 9 | Donkey F(ab')2 Anti-Sheep IgG H&L (PE) preadsorbed | abcam | ab7009 |
| 10 | F(ab')2-Goat anti-Mouse IgG (H+L) Secondary Antibody, PE, eBioscience™ | Invitrogen | 12-4010-82 |
| 11 | Goat anti-Rabbit IgG (H+L) Cross-Adsorbed Secondary Antibody, PE | Invitrogen | P-2771MP |

**Table S2.** Primers used for genotyping.

| No | Target | Sequence | Direction | Explanation |
|---|---|---|---|---|
| 1 | *ACTB* | GAAGATCAAGGTGAGTGCCC | Forward | For amplifying target loci used in TIDE |
| 2 | | GGATTGTGATGGCTGACCAC | Reverse | |
| 3 | | GTCCCCTTCTCCTTCCAGAT | Forward | For Sanger sequencing of PCR amplicons used in TIDE |
| 4 | ACTB-mEGFP 5' junction | CTGGCACCACACCTTCTACA | Forward | For PCR verification of the ACTB–mEGFP 5' junction to confirm knock-in |
| 5 | | TCGCCCTTGCTCACCATTC | Reverse | |
| 6 | ACTB-mEGFP 3' junction | GACGTAAACGGCCACAAGTT | Forward | For PCR verification of the ACTB–mEGFP 3' junction to confirm knock-in |
| 7 | | TTTGCGCTGACAGTTCCATTT | Reverse | |
| 8 | *UBB* | CTGTGCCTTCACTCACAGGT | Forward | For amplifying target loci used in TIDE |
| 9 | | CTGGACAAAGGACACAGTTCG | Reverse | |
| 10 | | AGTTCGTACGCAGTGTTTAAC | Forward | For Sanger sequencing of PCR amplicons used in TIDE |
| 11 | UBB-mScarlet 5' junction | TCAGAAGCGACTGGCATTGT | Forward | For PCR verification of the ACTB–EGFP 5' junction to confirm knock-in |
| 12 | | GCCTCTGTGGAGTCCATGTC | Reverse | |
| 13 | ACTB-mScarlet 3' junction | TACCGAACGGCTTTATCCCG | Forward | For PCR verification of the ACTB–EGFP 3' junction to confirm knock-in |
| 14 | | TCGCCCTTTCCTTCTACCAC | Reverse | |

**Table S3.** Antibodies used for Simple Western analysis.

| No | Name of the antibody | Manufacturer | Cat# |
|---|---|---|---|
| 1 | Human/Mouse/Rat Phospho-ATM (S1981) Antibody | R&D Systems | AF1655 |
| 2 | Phospho-p53 (Ser15) Antibody | Cell Signaling Technology | 9284 |
| 3 | Anti-Caspase-3 antibody [ABM1C12] | abcam | ab208161 |
| 4 | Caspase-7 Antibody | Cell Signaling Technology | 9492 |
| 5 | Phospho-DNA-PK (Thr2609) Polyclonal Antibody | Invitrogen | PA5-105749 |
| 6 | Anti-gamma Actin antibody [2A3] | abcam | ab123034 |
